# Earthworms and Cadmium - heavy metal resistant gut bacteria as indicators for heavy metal pollution in soils?

**DOI:** 10.1101/295444

**Authors:** Maja Šrut, Sebastian Menke, Martina Höckner, Simone Sommer

## Abstract

Preservation of the soil resources stability is of paramount importance for the ecosystem, particularly in the current era of environmental change, which presents a severe pollution burden (e.g. by heavy metals) to soil ecosystems and its fauna. Gut microbiomes are becoming recognized as important players in organism health, with comprehension of their perturbations in the polluted environment offering new insights into the nature and extent of heavy metal effects on the health of soil biota. Our aim was therefore to evaluate the effect of environmentally relevant heavy metal concentrations of cadmium (Cd) on the earthworm gut microbiota. Cd exposure led to perturbations of several heavy metal resistant taxa as well as taxa able to bind heavy metals, revealing the potential of the earthworm-microbiota system in overcoming human-caused heavy metal pollution. An ‘indicator species analysis’ linked bacterial genera *Paenibacillus* and *Flavobacterium*, and members of the order Actinomycetales with Cd treatment, suggesting the possible use of these bacterial taxa as biomarkers of exposure for Cd stressed soils. The results of this study will be essential to understanding of the soil fauna health, under anthropogenic disturbance, and will have implications for environmental monitoring and protection of soil resources.

**Importance:** Soil heavy metal pollution presents a severe burden for soil invertebrates and can have impact on their health, which in turn reflects on the health of the entire ecosystem. Gut microbiome is recognized as a central driver of the host health and its shifts can have severe consequences for the host. In this study we investigated the impact of cadmium (Cd) on earthworm gut microbiota, in a controlled experiment using cutting edge next generation sequencing and state of the art bioinformatics tools. The significance of this study is in identifying the gut bacterial taxa which are indicators for Cd treatment and are potential biomarkers of exposure to Cd. Therefore, this study contributes to develop efficient measures to qualify environmental pollution and to protect fragile soil resources and ultimately human health.

## Introduction

Environmental pollution by heavy metals poses a severe risk for soil ecosystems. Cadmium (Cd), one of the main heavy metal soil pollutants, appears in soil ecosystems as a side product of the metal and mining industry, from usage of fertilizers containing Cd and through air deposition (1). Heavy metals, including Cd, are linked to various toxic effects in exposed organisms, such as the induction of oxidative stress, DNA damage and carcinogenesis, and effects on the immune system (2). Metal-polluted soils also strongly influence soil microbiota and the microbiota of soil organisms. Several studies have shown a decrease in microbe biodiversity in metal-polluted soils (3–5). For instance, Cd inhibits microbial reproduction in soil (6), and Cd, copper (Cu), nickel (Ni), lead (Pb), chromium (Cr), arsen (As) and zinc (Zn) have been found to decrease the biomass, species richness and activity of microbial communities in forest and arable soils (7–10). Microbial communities in heavily polluted soils have been hypothesized to be reduced to only 1 % of that usually observed in pristine soils (11).

Earthworms are dominant members of the soil macrofauna and are essential species in soil. Because of their burrowing activities and casting, earthworms alter the structure of the soil and thus contribute to the cycling of nutrients and to geochemical soil processes (12). Earthworms carry out soil biological regulation by mixing organic matter and mineral particles inside their gut. Further, they maintain the soil structure, and regulate the water content and availability of nutrients for plants (13). Thereby, they also have an impact on soil microbial properties, an aspect that is crucial for the stability of soil ecosystems (14). The earthworm gut represents a unique ecological niche with stable conditions, in contrast to the surrounding soil conditions (15). Although earthworm gut microbiota have been previously thought to resemble the bacterial diversity present in its environment, this premise has been discarded with the description of the core earthworm microbiota (14, 16). Bacterial taxa present in the earthworm gut represent those found in the environment (diet, soil) only to a certain extent. This results from the filtering and specific milieu in their gut, such as a neutral pH, aerobic conditions, constant moisture, and high amount of carbon substrate (14). By ingesting soil bacteria during feeding and burrowing, earthworms act as biological filters for soil microbiota, containing specific microorganism groups, especially anaerobes (16–18). Additionally, the earthworm gut has the ability to discriminate between beneficial and harmful bacteria. For instance, some bacteria show tolerance to the antimicrobial activity of digestive fluids derived from the earthworm gut (19, 20). The maintained earthworm gut harbours a microbial community that is involved in food metabolism, thereby establishing bioavailable vitamins and nutrients, and pathogen protection (20–22). Bacteria involved in these processes fall into one of the four physiological groups: plant growth promoters, free-living nitrogen fixers, biocides and phosphate solubilizers (13). Interestingly, the microbiota of different earthworm species harbour comparable proportions of bacterial phyla. For instance, the core microbiota of *Lumbricus rubellus* comprises of Actinobacteria and Proteobacteria, with lower abundances of Bacteroidetes and Acidobacteria (14). In *Eisenia andrei* the majority of the gut microbiota belongs to the phyla Actinobacteria and Proteobacteria (16). In *E. fetida* and *Perionyx excavates*, the majority of bacteria belongs to the Proteobacteria and Firmicutes, followed by unclassified bacteria (23).

The gut microbiota interacts with ingested heavy metals, acts as an important mediator of metal bioavailability and toxicity and, furthermore, behaves as a barrier for the uptake of heavy metals (24). For instance, germ-free mice absorb significantly higher levels of Cd and Pb compared with control mice and as a result show different expression profiles of genes involved in metal detoxification and transport (25). Several metals have been linked to alterations in the gut microbiota in vertebrate species, such as Cd (26–29), Ni (30), As and iron (Fe) (31). In earthworms, environmental exposure to As and Fe causes shifts in the composition of the *L. rubellus* microbiota (14). A healthy and stable gut microbiota plays an important role in host health and changes in microbiota composition, known as dysbiosis, can have severe consequences. For instance, the microbiota and the host immunity are well known to be in continuous interaction, thereby modifying the host immune response (32). Gut microbiota changes can lead to a misbalance between the host immune system and the gut microbiota, can trigger immune responses and can lead to immunological diseases (2). Germ free mice have deficiencies in the development of lymphoid tissue, reduced innate and adaptive immune response and are more susceptible to infections (33). On the other hand, the immune response of the host can have an impact on the intestinal microbial composition and its function (34). The immunological control of the gut microbiota is critical for maintaining intestinal homeostasis and involves a variety of innate and adaptive components (34). Therefore, the changes in the host gut microbiota can serve as a first indication of potential misbalance in an organism.

Recent studies imply that the changes in gut microbiota structure and function can be used to evaluate the biological effects of pollutants and to define specific microbiota communities as biomarkers of exposure (24, 35). Therefore, an understanding of microbiota changes attributable to exposure to common pollutants in key organisms and the definition of chemically specific microbiota alterations should improve our comprehension of the nature and extent of pollution effects and enable better monitoring and preventative measures in the various ecosystems challenged with anthropogenic pollution.

Our aim has been 1) to investigate microbiota shifts in the earthworm *L. terrestris* exposed to various concentrations of the heavy metal Cd and 2) to identify bacterial indicator sequence variants involved in the first response of the earthworm gut microbiota to Cd stress in order to propose potential microbiota biomarkers of exposure.

## Results

### Sequencing and assignment of SVs

High-throughput amplicon sequencing of the V4-region of the bacterial 16S rRNA gene from soil, manure and earthworm faecal samples resulted in a total of 14,815,785 reads. Nine samples dropped out because of low sequencing depth (max. sequence depth: 7). After quality filtering, 10,247,539 reads remained for the assignment of SVs for 178 samples (min: 30,932; max: 91,298; mean: 57,570; median: 57,710). These reads were assigned to 4,787 SVs (min: 1.0; max: 814; mean: 2,140; median: 52).

### Soil and manure microbiota

All soil samples had higher α-diversities than earthworm gut microbiota samples and manure samples (Figure 2). Soil sampled before earthworm introductions (Soil_acc) had a slightly higher α-diversity than soil after earthworm acclimation (Soil_Cd-0_pre), but this effect was not significant (Kruskall-Wallis: H = 1.19, *p* = 0.275, Figure 2). Interestingly, the control soil collected after earthworm exposure (Soil_Cd-0_post) had a lower, albeit non-significant (Kruskall-Wallis: H = 3.0, *p* = 0.083) α-diversity compared with the Cd-exposed soil (Soil_Cd-10/50_post), indicating that Cd treatment does not cause further decrease of the soil microbiota diversity, at least in the short term.

**Figure 2.**
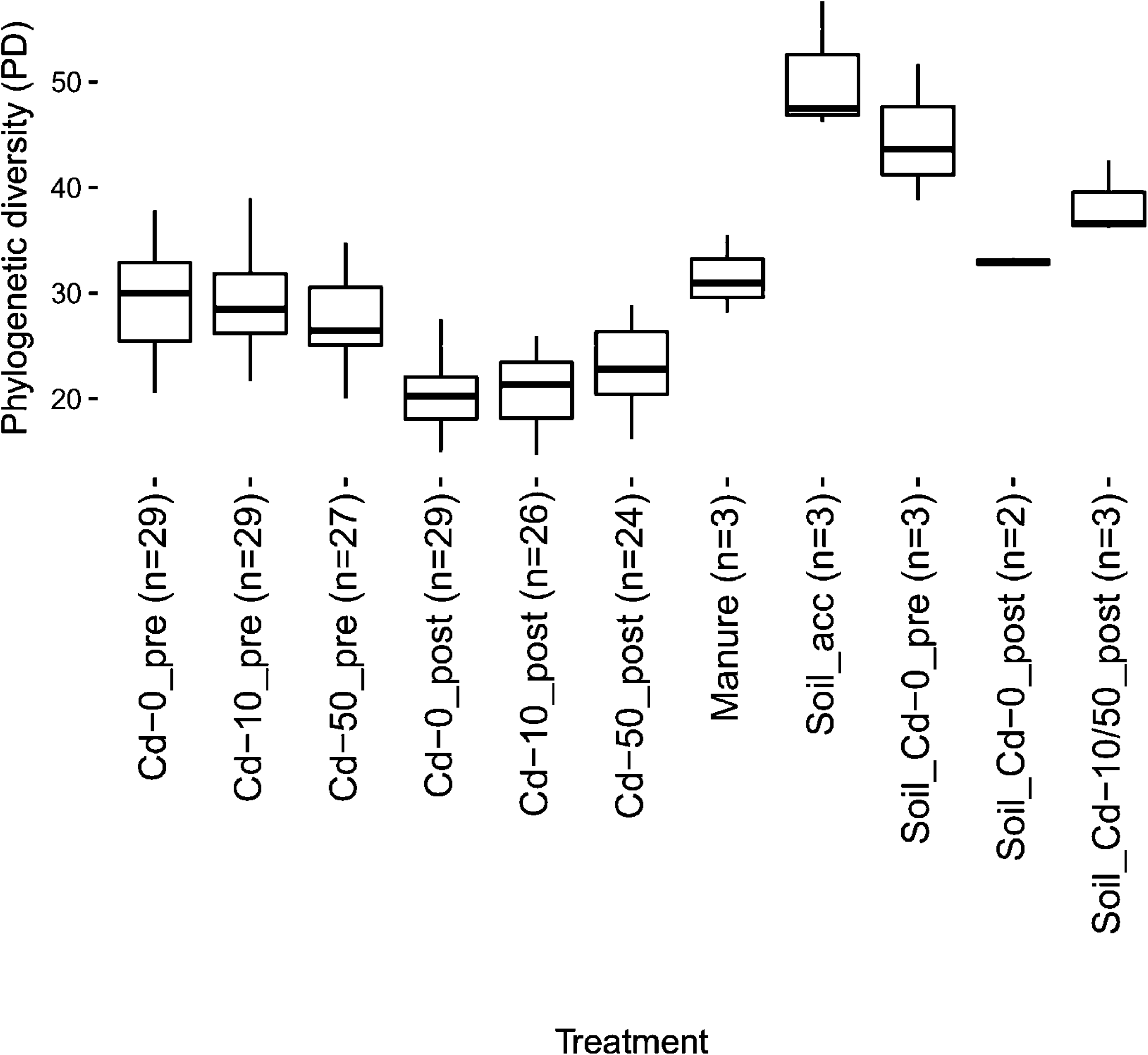
α diversity boxplots representing Faith’s Phylogenetic Diversity (PD) between treatment groups, soil and manure samples. Soil and manure samples have a higher PD compared with earthworm gut microbiota, independent of treatment group.

Neither soil nor manure samples possessed the phylum Tenericutes at proportions seen in the earthworm gut microbiota (Figure 3). ‘Soil_acc’ had a significantly higher abundance of Firmicutes and a lower abundance of Proteobacteria and Bacteroidetes compared with ‘Soil_Cd-0_pre’ (Figure 3, Table S1, supporting information (SI)). After earthworm acclimation in experimental soil (‘Soil_Cd-0_pre’, ‘Soil_Cd-0_post’), the proportion of beta and gammaproteobacteria increased in comparison with that in ‘Soil_acc’, whereas the abundance of alphaproteobacteria remained the same (Table 1, Table S2 (SI)).

**Table 1.**
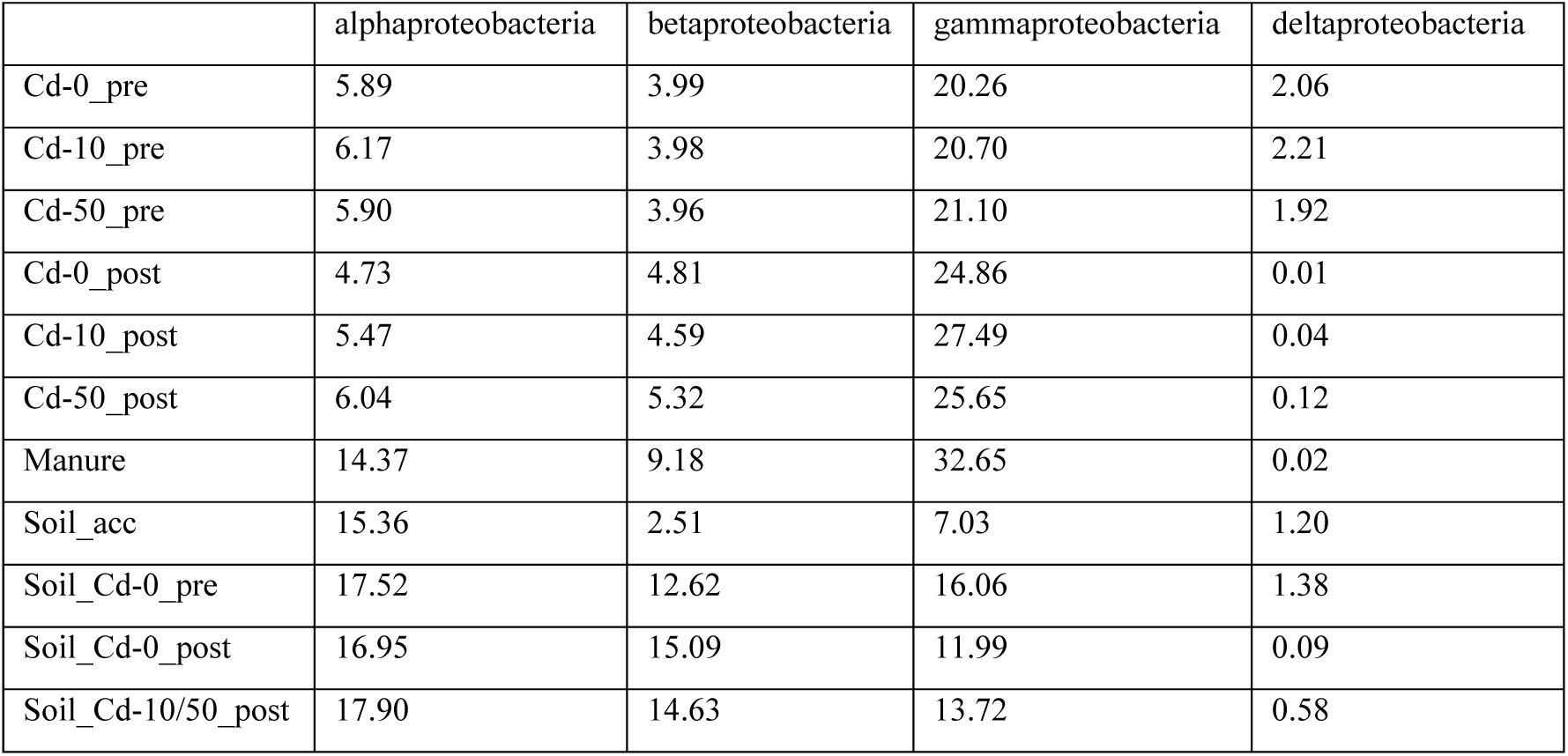
Relative proportion (%) of various Proteobacteria classes in treatment groups, soil and manure samples.

**Figure 3.**
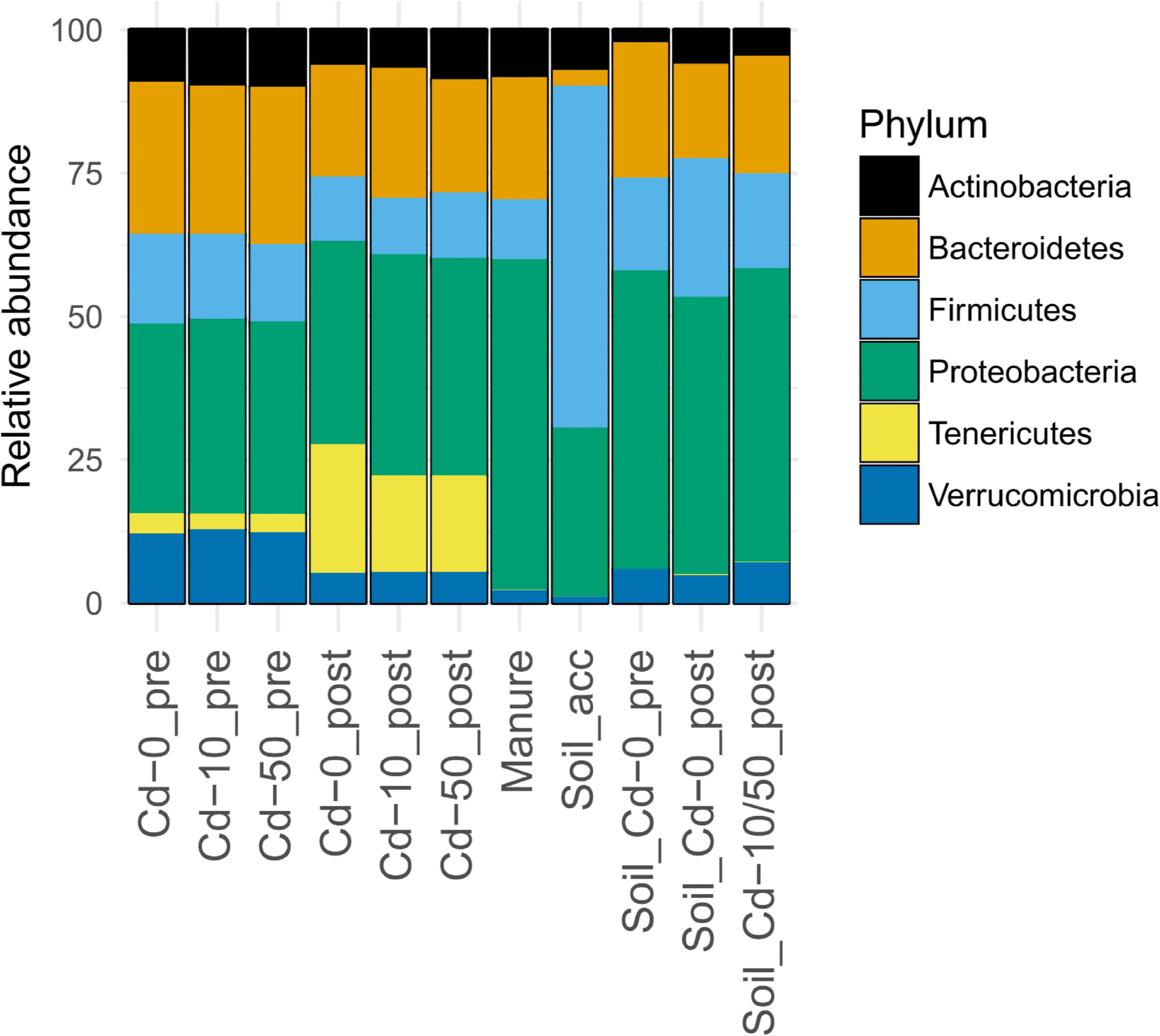
Proportions (%) of the most abundant phyla in the various earthworm treatment groups and in the soil and manure samples. Proteobacteria was the bacterial phylum with the highest proportion in all groups, other than for Soil_acc in which the phylum Firmicutes predominated.

### Earthworm gut microbiota changes attributable to Cd treatment

We observed significant differences in α diversity in the control groups between the first and second sampling (Figure 2). This indicates that the acclimation period of one month was not sufficient to stabilize earthworm gut microbiota. Thus, subsequent analyses focused on the comparisons between ‘Cd-0_post’, ‘Cd-10_post’, ‘Cd-50_post’, manure and soil samples.

Our results revealed that Cd exposure caused perturbations in earthworm gut microbiota composition. A significant increase in α diversity could be observed for treatment ‘Cd-50_post’ compared with ‘Cd-0_post’ (Kruskall-Wallis: H = 5.07, *p* = 0.02) (Figure 2). Furthermore, weighted UniFrac distances were significantly different between treatment groups (F_2, 76_ = 3.30, *p*= 0.006), which is also evident from the PCoA plot (Figure 4). Moreover, the model containing ‘treatment group’ as explanatory variable explained more variation than the model without (Figure 5, Table S3 (SI)).

**Figure 4.**
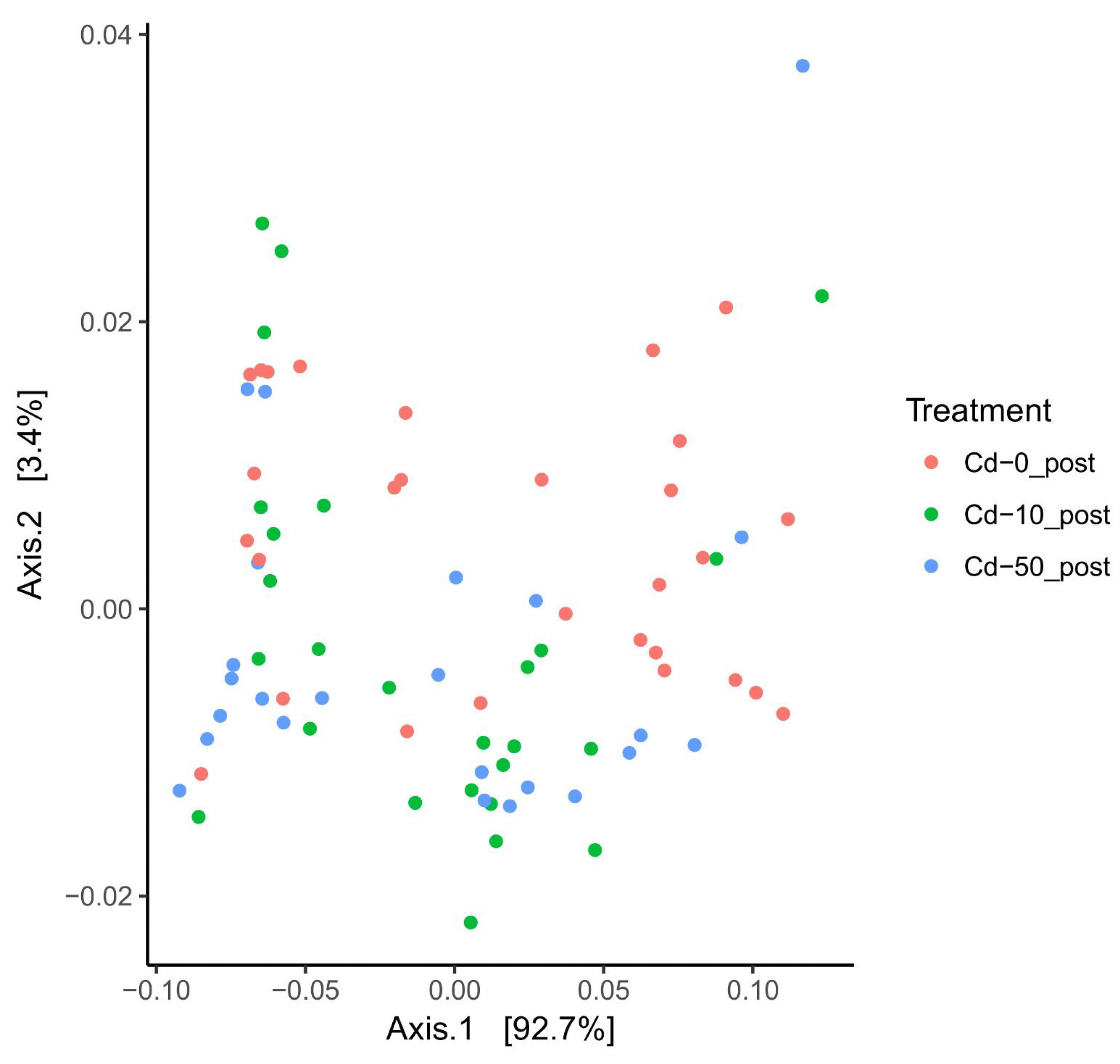
PCoA plot based on the weighted UniFrac matrix revealed the clustering according to the treatment groups relative to the control group.

**Figure 5.**
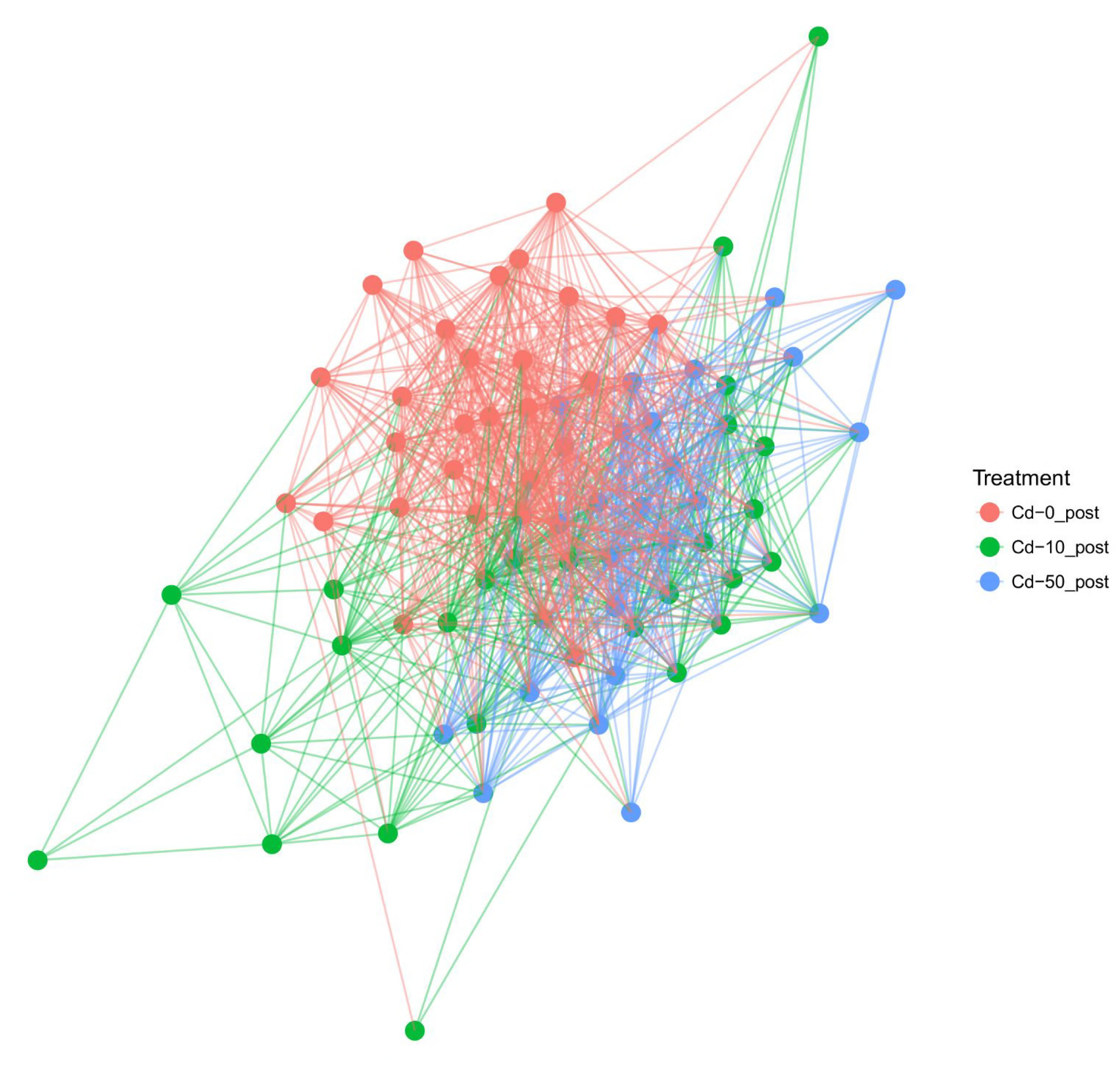
A network graph showing the connectedness between earthworm gut microbiota according to treatment group.

*L. terrestris* microbiota in the combined pre-exposure groups ‘Cd-0_pre’, ‘Cd-10_pre’ and ‘Cd-50_pre’ mostly consisted of Proteobacteria (32.75 %), Bacteroidetes (26.02 %), Firmicutes (14.29 %), Tenericutes (3.07 %), Verrucomicrobia (12.15 %) and Actinobacteria (9.28 %). In the control samples ‘Cd-0_post’, these relative frequencies differed as follows: Proteobacteria (34.53 %), Bacteroidetes (18.90 %), Firmicutes (10.94 %), Tenericutes (21.72 %), Verrucomicrobia (5.08 %) and Actinobacteria (5.96 %) (Figure 3). A significantly higher abundance of Tenericutes and a lower abundance of Verucromicrobia were evident between all groups before exposure (‘Cd-0_pre’, ‘Cd-10_pre’, ‘Cd-50_pre’) and the control group after the exposure (‘Cd-0_post’) (Table S1 (SI)). Among Proteobacteria in the earthworm gut, the most abundant were gammaproteobacteria, followed by alphaproteobacteria and betaproteobacteria, whereas deltaproteobacteria were present in low numbers. The abundance of alpha- and betaproteobacteria in the earthworm gut was significantly lower in comparison with that in the soil samples. In contrast, the abundance of gammaproteobacteria was higher in the gut samples, although not significantly (Table 1, Table S2 (SI)). Cd treatment at the concentration of 50 mg/kg (‘Cd-50_post’) caused significant alteration in the proportion of Actinobacteria in comparison with the ‘Cd-0_post’ group (Figure 3, Table S1 (SI)).

### Cd-sensitive and Cd-resistant taxa

A heatmap visualizing all of the balances obtained through Gneiss analysis revealed differences between the treatment groups (Figure S1 (SI)). In order to identify whether earthworm gut microbial SVs differed significantly between non-polluted and Cd-polluted groups, we compared the abundance of SVs by using DESeq2 analysis. Out of the SVs that could be assigned to a bacterial genus in ‘Cd-10_post’, 23 SVs increased, whereas 6 SVs decreased in abundance in relation to ‘Cd-0_post’ (Figure 6a). A strong increase was observed for SVs of the genera *Flavobacterium* (up to 23.51 fold), *Sanguibacter* (23.24 fold), *Dermacococcus* (7.34 fold), *Paenibacillus* (6.42 fold), and *Fluviicola* (5.57 fold). A strong negative response was observed for SVs of the genera *Cryocola* (-6.28 fold) and *Perlucidibaca* (-24.38 fold). In ‘Cd-50_post’, 33 SVs increased, whereas 15 SVs decreased in abundance in relation to ‘Cd-0_post’ (Figure 6b). A strong increase was observed for SVs of the genera *Luteolibacter* (up to 23.65 fold), *Salinbacterium* (23.18 fold), *Cellulomonas* (9.71 fold), *Rhatayibacter* (9.42 fold), *Dermacoccus* (7.93 fold), *Nocardioides* (6.88 fold), *Paenibacillus* (6.24 fold), *Candidatus xiphinematobacter* (5.74 fold), *Flavobacterium* (5.52 fold), *Chryseobacterium* (5.21 fold) and *Agrobacterium* (4.69 fold). Similar to ‘Cd-10_post’, a negative response was observed for SVs of the genera *Cryocola* (-2.63 fold) and *Perlucidibaca* (-4.76 fold).

**Figure 6.**
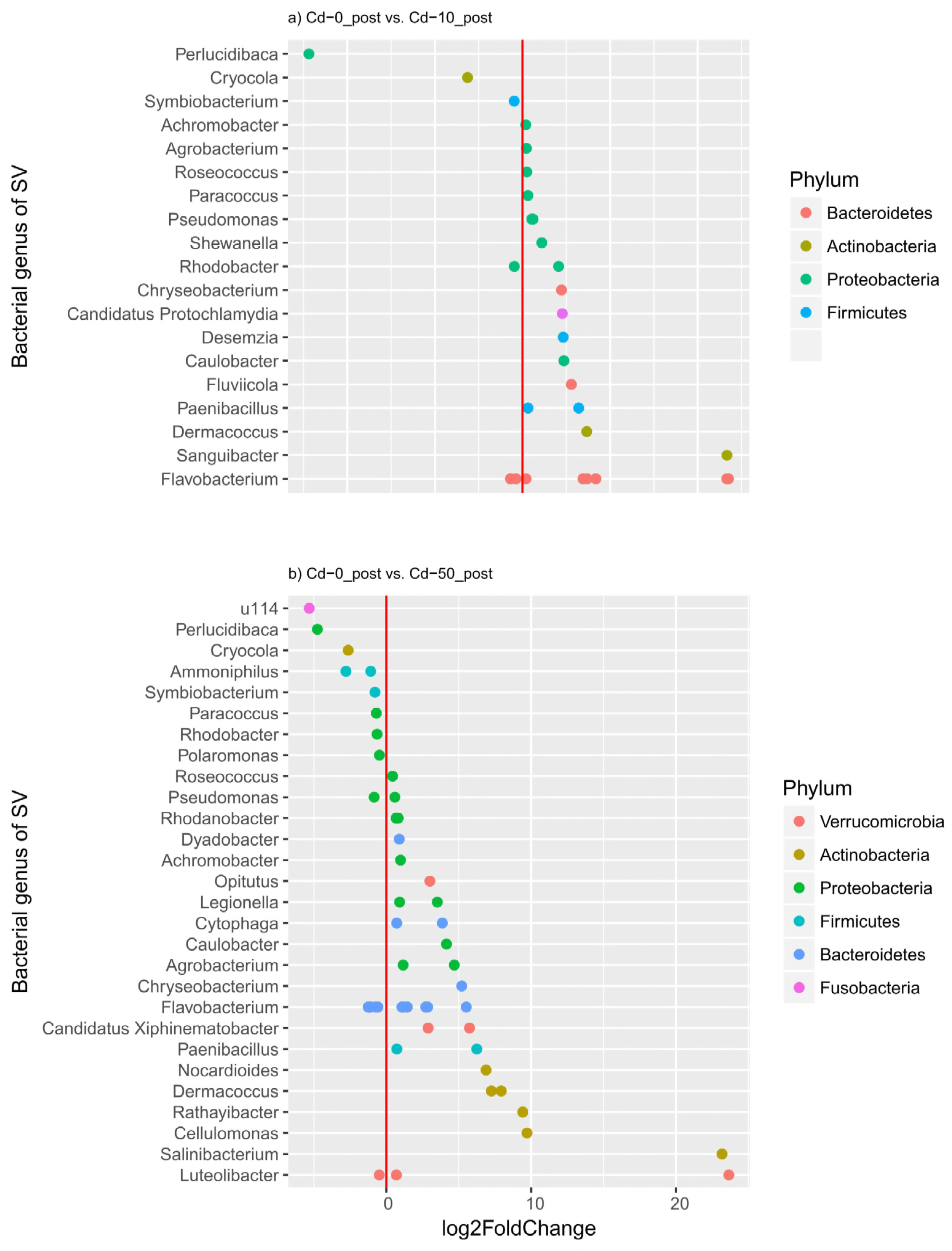
Results of DESeq2 analyses showing log2-fold changes of bacterial taxa for comparisons between Cd-0_post and Cd-10_post and between Cd-0_post and Cd-50_post.

### Unique bacterial indicator taxa associated with Cd treatment

Indicator species analysis was employed in order to find predictive patterns of Cd exposure. For the control treatment (‘Cd-0_post’), we identified a total of 13 indicator SVs, which included mostly members of the orders Legionellales and Bacillales. 23 and 57 indicator SVs were linked to the 10 and 50 mg/kg Cd treatment, respectively. Many of the Cd associated SVs overlapped between the 10 and 50 mg/kg treatments including mostly members of the genera *Paenibacillus* and *Flavobacterium*. Other overlaps included the genera *Candidatus*, *Chthoniobacter* and *Rathayibacter* and the bacterial families Streptomycetaceae, Verrucomicrobiaceae, Comamonadaceae and Chthoniobacteraceae (Figure 7). In the ‘Cd-50_post’ treatment group many indicator taxa belonged to the order Actinomycetales, including the families Microbacteriaceae (genera *Rathayibacter* and *Agromyces*), Nocardioidaceae (genera *Nocardioides* and *Pimelobacter*), Nocardiaceae (genus *Rhodococcus*), Dermacoccaceae (genus *Dermacoccus*), Micrococcaceae and Geodermatophilaceae (Figure 7).

**Figure 7.**
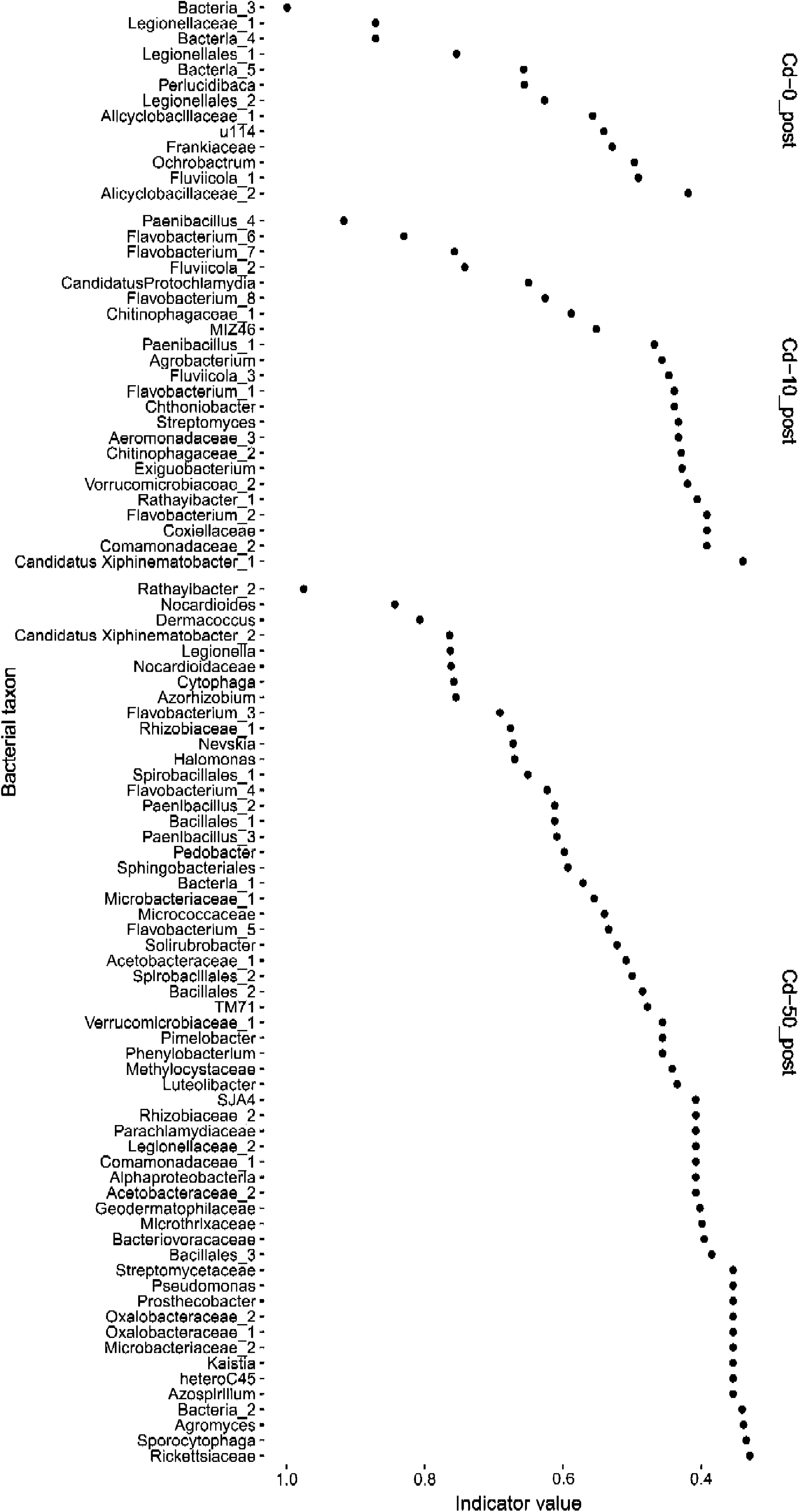
Indicator species that significantly characterize treatment groups after earthworm exposure. An indicator value of 1 indicates that a certain SV is present in all individuals of a group. An indicator value less than 1 means that the SV is either present in more than one group or that it is not present in all members of this group.

## Discussion

### Earthworm gut microbiota differs from soil and manure microbiota

The α diversity of earthworm gut samples was lower in comparison with that of soil and manure samples and, as expected, did not fully represent the microbiota community found in the experimental soil or manure used for feeding. This is in accordance with previous findings showing that the microbiota associated with the earthworm gut has a reduced level of diversity and richness compared with the surrounding soil environment (14, 36). The proportion of Proteobacteria and Bacteroidetes was significantly lower, whereas the proportion of Firmicutes was significantly higher in ‘Soil_acc’, in comparison with ‘Soil_Cd-0_pre’ and ‘Soil_Cd-0_post’, indicating the profound impact of earthworms on the soil microbial community. Through soil ingestion, earthworms modify microbial composition and activity by depositing casts in the soil in which microbes can either flourish or decline (37). As previously shown, earthworm casts have a similar bacterial composition to the surrounding soil but differ in their proportions (38). Manure had a higher proportion of Proteobacteria in comparison with ‘Soil_acc’ and, thus, the combined influence of the food source and earthworm activity might have led to the observed changes in the soil microbial composition prior to and after earthworm introduction.

The proportion of certain bacterial taxa differed between the earthworm gut and soil samples after the acclimation period. For instance, gammaproteobacteria were more abundant in gut samples in comparison with ‘Soil_Cd-0_pre’ and ‘Soil_Cd-0_post’, whereas alpha- and betaproteobacteria were less abundant. Similarly, in another study, alpha-, beta- and gammaproteobacteria declined in the *L. terrestris* gut in comparison with their values in the soil (39). Additionally, gammaproteobacteria have also been reported to increase in the earthworm gut and cast samples in comparison with the surrounding soil (38).

The phylum Tenericutes, consisting mostly of undetermined orders of the class Mollicutes, was missing from soil and manure samples but was abundant in the earthworm gut. The increase of Mollicutes in the gut after the exposure period (2 months in experimental soil) in comparison with the abundance before exposure (1 month in the experimental soil) indicates that this phylum is strongly linked to the earthworm gut and is not derived from either soil or food. Nevertheless, we cannot totally discard the premise that it might be derived from the soil in which earthworms were originally grown by the breeder. However, bacteria from the Mollicutes class have previously been described in the gut and coelomic fluid of Lumbricidae earthworms, although no conclusion could be made as to whether the earthworm gut is their specific habitat (15, 40).

### Dose-dependent effect of Cd on microbiota of earthworm gut

The most abundant phylum in the *L. terrestris* gut samples was Proteobacteria, followed by Bacteroidetes, Firmicutes, Tenericutes, Verrucomicrobia and Actinobacteria. This is similar to the previously described *L. terrestris* gut microbiota composition consisting predominantly of Proteobacteria, followed by Bacteroidetes and with lower abundances Acidobacteria, Planctomycetes, Verrucomicrobia and Actinobacteria (15). Proteobacteria also seem to be the most abundant phyla for other earthworm species. For instance, in *L. rubellus*, Proteobacteria comprised more than 50 % of the gut microbiota, followed by Actinobacteria, Bacteroidetes and Acidobacteria, whereas Firmicutes, Chloroflexi and Cyanobacteria appeared in lower proportions (14). In *E. andrei* the majority of microbiota belonged to the phyla Proteobacteria and Actinobacteria (16). Similarly, in *E. fetida* and *P. excavates*, the majority of the observed microorganisms were Proteobacteria, followed by Firmicutes and unclassified bacteria (23).

In our study we observed a significant increase in the proportion of Actinobacteria after 50 mg/kg Cd treatment in comparison with the control. Similarly, in mice orally treated with Cd for 8 weeks, the proportion of Actinobacteria in the cecal content changed significantly at a concentration of 100 ppm compared with the control (26). Heavy metals have previously been linked to the alterations of the gut microbiota community in various vertebrate and invertebrate organisms, including earthworms (14, 26, 30, 31). For Cd, the effects on the microbiota community have been described only in mice and fish. For instance, sub-chronic exposure of mice to a low dose of Cd caused a decrease in the abundance of Firmicutes and gammaproteobacteria and an increase in the abundance of Bacteroidetes in the gut and this was linked to hepatic inflammation and dysregulation of energy metabolism (28). Similarly, a lowdose Cd exposure in early mouse life stages caused an increase in Bacteroidetes and decrease in Firmicutes (41). An increase in the abundance of Bacteroidetes bacteria was also observed in Nile tilapia treated with Cd (29).

Bacteria of the phylum Tenericutes and particularly the class Mollicutes decreased in abundance following Cd treatment at both concentrations compared with the control. Although this change was not significant, a similar pattern was previously described for Tenericutes in amphibian gut microbiota exposed to long-term heavy metal pollution (24).

### Cd-sensitive and Cd-resistant taxa were determined among earthworm gut microbiota

Many of the genera that showed a Cd-dependent increase in abundance belonged to the phylum Actinobacteria. Among this phylum, 6 genera showed a particularly strong increase, namely *Dermacoccus*, *Rathayibacter, Nocardioides, Sanguibacter, Salinbacterium* and *Cellulomonas*. Increasing numbers of *Rathayibacter*, *Nocardioides* and *Cellulomonas* attributable to Cd treatment can be explained by their reported resistance to heavy metal toxicity. Metal tolerant strains of *Rathayibacter* have been found in soils with historic heavy metal contamination (42, 43). Similarly, *Nocardioides* has a high adaptive potential to survive and maintain metabolic activity under extreme conditions (heavy metals, UV, nuclear radiation) (44). *Cellulomonas sp.* has been reported to be highly resistant to heavy metals and the *C. hominis* N2 strain has the ability to produce a biosurfactant, that might help to overcome heavy metal toxicity (45, 46). The abundance of several phyla with a reported resistance to heavy metals increased in our study, namely *Paenibacillus, Flavobacterium, Chryseobacterium* and *Agrobacterium.* Some of the *Paenibacillus* strains can survive high heavy metal concentrations and their possible role in heavy metal decontamination makes them possible candidates for heavy metal bioremediation (47, 48). Likewise, *Flavobacterium* species show resistance to some heavy metals (Cd, Cu) and their abundance in Nile tilapia gut upon exposure to Cd was significantly increased (29, 49). *Chryseobacterium solincola* can grow despite the elevated presence of highly toxic metals and has been suggested as a candidate for the situ bioremediation of heavy metals in aqueous or soil systems (50). *Agrobacterium tumefaciens* is able to survive in regions containing high levels of heavy metals and possesses various transporters involved in Ni, Cu, Zn and Cr resistance (51, 52). SVs of the genera *Cryocola* and *Perlucidibaca* significantly decreased in abundance in both Cd treatments. This is in contrast to previous findings reporting a positive correlation of *Perlucidibaca* bacteria with Cd content in samples of polluted river sludge (53).

### Unique bacterial indicators as potential biomarkers of exposure to Cd

In order to reveal a unique microbiota footprint in the earthworm gut attributable to Cd exposure, we have identified indicator SVs by using indicator species analysis (54). This analysis has previously been used to characterise microbial taxa in the human gut related to diverse diets (55), in order to reveal microbial biomarkers of disease (56), to define microbial taxa associated with various environments (57) and, most recently, to identify biomarkers of exposure to chemical pollution in fish (35). The authors of the last-mentioned study suggest that the characterization of indicator SVs of the microbiota exposed to environmental chemicals may be useful in environmental and health monitoring. Many of the indicator SVs associated with Cd exposure in our study belonged to the genera *Paenibacillus* and *Flavobacterium*. These genera also showed increasing numbers following Cd treatment in the DESeq analysis. Members of these genera are soil-derived denitrifiers or dissimilatory nitrate reducers in the earthworm gut and contribute to the nitrous oxide emissions and help in organic matter digestion by providing hydrolytic enzymes (58). Many of the other Cd-associated indicator SVs belonged to the order Actinomycetales, including genera *Dermacoccus*, *Rathayibacter* and *Nocardioides*, which were also significantly associated with Cd treatment in the DESeq2 analysis. Other indicator taxa from this order were genera *Streptomyces*, *Agromyces*, *Pimelobacter* and *Rhodococcus* and families Micrococcaceae and Geodermatophilaceae. Actinomycetales genera *Streptomyces*, *Rathayibacter*, *Nocardioides*, *Agromyces* and *Rhodococcus* have been associated with the soil and earthworm cast and gut. The genus *Streptomyces* has antimicrobial activity and enzyme production capability in the earthworm gut (59). Furthermore, *Streptomyces caeruleus* has been found to help earthworms in organic matter metabolism and plant material decomposition (60). *Rathayibacter* is a known genus of soil bacteria and contains plant pathogens (61). *Nocardioides* has been associated with the cast of the earthworm *E. andrei* (62) and *Agromyces* bacteria and the *Rhodococcus* strain has been isolated from the earthworm gut (63, 64). Evidence has been presented for the metal tolerance for these Actinomycetales indicator taxa. Streptomyces strains bioaccumulate various heavy metals (Zn, Cu, Cd, Cr, Ni, Sr, U) and are able to remove Zn from soil (65). The species *Agromyces aureus* was isolated from a zinc/lead mine and is described as a heavy-metal-resistant bacteria, with a resistance of up to 1 mM of Cd (66). *Rhodococcus opacus* has been described to be able to remove heavy metals from the media by active bioaccumulation and to possess tolerance to a range of heavy metals, including Cd (67, 68). Various species of the Micrococcaceae family have been reported as being resistant to high levels of heavy metals, including Cd (69), and members of the Geodermatophilaceae family have been determined to be tolerant to a range of stresses, including heavy metals (70). Among other indicator SVs, those that were associated with both low and high Cd treatments included the genera *Candidatus* and *Chthoniobacter* and the families Verrucomicrobiaceae, Comamonadaceae and Chthoniobacteraceae. The genus *Chthoniobacter* and the family Verrucomicrobiaceae have been found in soil ecosystems and *Chthoniobacter* is probably involved in the breakdown of organic carbon in soil (71, 72). Members of the phylum Verrucomicrobia have been detected in mercury contaminated sites (73). Comamonadaceae of the phylum Proteobacteria have been isolated from the digestive tracts of diverse earthworm species (74) and some bacterial strains of this family have been associated with arsenic-contaminated earthworm microbiotas and are described to carry arsenic-resistance genes (14, 75). Defined indicator SVs for both low and high Cd treatments might serve as a catalogue of biomarkers of exposure to Cd and be used for future biomonitoring programmes in which earthworms could be used as an appropriate model to describe bacterial indicator taxa for human-caused environmental pollutants.

Because of the high extent of chemical exposure to soil organisms and the overall role of the gut microbiota for organism health, the description and comprehension of the effects of chemical exposure on the microbiota are of high importance. The overall results of this study should help us to understand the dynamics and the effects of heavy metal pollution on the microbiota of key soil invertebrates and thereby to contribute to measures for preserving environmental health.

## Materials and methods

### Earthworm exposure and sampling

All *L. terrestris* specimens and soil (80 g dry soil per earthworm) used for earthworm exposure (mixture of peat and humus) were obtained from the company Wurmwelten/Germany (www.wurmwelten.de). Ninety adult earthworms originating from a single population were reared at 15°C in a 12h/12h light/dark cycle in heat-treated soil (120°C, 12 h) with a soil water content of 50% and fed weekly with horse manure (1.2 g manure per individual) for 4 weeks prior to the start of the experiments (acclimation period). Following this acclimation period, earthworms were divided into three groups of 30 individuals that were exposed either to control soil or to 10 mg/kg or 50 mg/kg CdCl_2_ for 4 weeks. The gut content of earthworms was sampled prior to and after exposure. Gut samples collected prior to exposure were labelled as ‘Cd-0_pre’, ‘Cd-10_pre’ and ‘Cd-50_pre’, whereas samples collected after the exposure are labelled ‘Cd-0_post’, ‘Cd-10_post’ and ‘Cd-50_post’. The experimental setup is schematically shown in Figure 1. Faecal matter was collected by placing individual earthworms overnight at 15°C in the dark on a sterile moist filter paper placed in a Petri dish. The released pellet was collected by using a sterile spatula, placed in a 1.5 ml tube, flash-frozen in liquid nitrogen and stored at -80°C until further processing. In addition, soil and manure samples were collected in order to investigate their microbiota communities. ‘Soil_acc’ refers to samples of freshly prepared experimental soil for acclimation before the introduction of earthworms, ‘Soil_Cd-0_pre’ refers to soil samples collected after 4 weeks of earthworm acclimation, ‘Soil_Cd-0_post’ refers to soil samples with the treatment ‘Cd-0_post’ and ‘Soil_Cd-10/50_post’ refers to soil samples from ‘Cd-10_post’ and ‘Cd-50_post’ treatments, which were grouped together because some of the replicas were not successfully sequenced.

**Figure 1.**
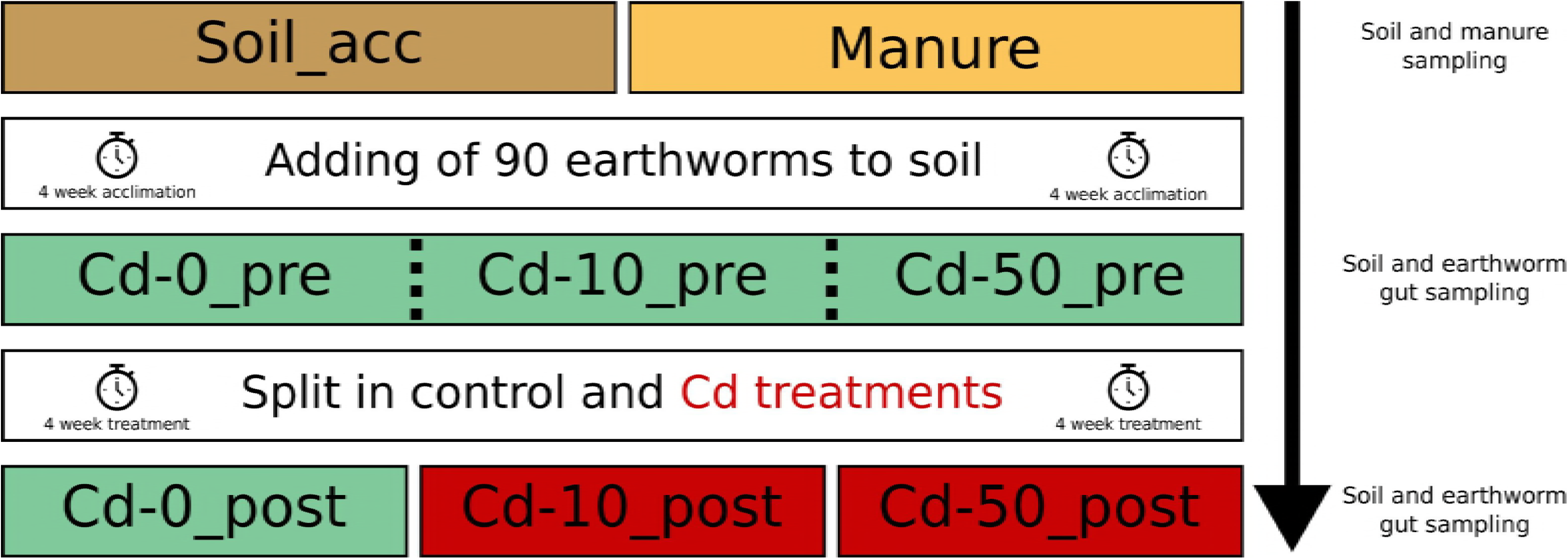
Schematic representation of the experimental setup. Earthworms were exposed to heavy metal (Cd) and their gut microbiota were sequenced before and after exposure to various Cd concentrations. The respective soil and manure microbiotas were also sequenced.

### DNA extraction, library preparation and sequencing

We used 200 mg earthworm gut content, soil and manure samples for DNA extraction by using the NucleoSpin 96 Soil kit (Macherey&Nagel). Mechanical lysis was performed for 2×2.5min at 50 Hz by using the Analytic Jena Homogenizer.

For library preparation, we applied the Fluidigm (Access Array^TM^ System for Illumina Sequencing Systems, Fluidigm Corporation) approach and chemistry for simultaneous PCR and barcoding. A 291-bp fragment of the hypervariable V4 region of the 16S rRNA gene was targeted by using the primers 515F (5′-GTGCCAGCMGCCGCGGTAA-3′) and 806R (5′-GGACTACHVGGGTWTCTAAT-3′) (76, 77). The primers were modified according to the Fluidigm protocol and were tagged with sequences (CS1 forward tag and CS2 reverse tag) that were complementary to the forward or reverse access array barcode primers for Illumina. The reaction was performed in 15 µl and consisted of 10 ng/µl template DNA, 1X FastStart HighFidelity Reaction buffer without MgCl_2_ (Roche), 4.5 nM MgCl_2_ (Roche), 5% DMSO (Roche), 200 μM of each PCR grade nucleotide (Roche), 0.05 U/μl FastStart High Fidelity Enzyme blend (Roche), 400 nM access array barcode primers for Illumina (Fluidigm), 200 nM target specific primers and 14% PCR certified water. PCR cycles were performed according to the Fluidigm protocol. Subsequently, the PCR samples were cleaned by using NucleoMag NGS Beads (Machery&Nagel) according to manufacturer’s recommendations and the cleaned libraries were quality checked with capillary electrophoresis on the Qiaxcel Advanced system (Qiagen) and quantified by using the Quant-iT™ PicoGreen ® kit (Invitrogen/Life Technologies). Samples were pooled with equal amounts of 150 ng DNA/sample. Finally, pooled samples were diluted to 8 nM in hybridization buffer and libraries were sequenced as paired-end run on Illumina ® MiSeq.

### Bioinformatic analyses with qiime2

Pre-processing of sequencing reads was carried out by using qiime2 (version 2017.10) ((78), https://qiime2.org) and its plugins. Specifically, we used the ‘demux’ plugin (https://github.com/qiime2/q2-demux) for the import of our demultiplexed paired-end sequencing reads and the creation of the ‘artifact’ file (i.e. qiime2 data format required for subsequent analyses). Further, we applied the ‘dada2’ plugin (79), by using the default parameter settings for quality filtering and chimera filtering, to trim primers (--p-trim-left-f 23, --p-trim-left-r 20), to truncate forward and reverse reads (--p-trunc-len-f 200, --p-trunc-len-r 200) and finally to collapse reads into representative sequences, the so-called sequence variants (SVs). We assigned taxonomy to these SVs against the Greengenes database (version 13_8) by using the ‘feature-classifier’ plugin (https://github.com/qiime2/q2-feature-classifier) with the ‘fit-classifier-sklearn’ method and produced taxa summary barplots (https://github.com/qiime2/q2-taxa) according to sample groupings.

In order to carry out diversity analyses, which are based on bacterial phylogeny, we produced a mid-point rooted bacterial phylogenetic tree by aligning SV with MAFFT (80), by removing non-informative positions in the alignment with the ‘mask’ command (https://github.com/qiime2/q2-alignment) and by using Fast Tree 2 (81) for tree construction. The ‘diversity’ plugin (https://github.com/qiime2/q2-diversity) was employed to calculate alpha (phylogenetic diversity, (82)) and beta diversities (weighted UniFrac, (83)) based on 30,000 sequences per earthworm microbiota. Significant differences between treatment groups were calculated with a PERMANOVA approach also included in qiime 2.

Finally, in order to be able to undertake further analyses in R (84), we exported the non-rarefied ‘feature-table’ (feature-table.biom), the bacterial phylogenetic tree (tree.nwk), and the taxonomy from qiime2 ‘artifacts’. We converted the ‘feature-table.biom’ file into a text file and then added the taxonomy in the last column and reconverted this text file into a ‘feature-table_tax.biom’ file both using the ‘biom convert’ command in qiime2 (http://biom-format.org, (85)).

### Statistical analyses in R

We imported and combined the ‘feature-table_tax.biom’ file, the bacterial phylogenetic tree, and a text file containing the metadata into R by using the phyloseq package (86). We produced the barplot including all treatments, the PCoA and the network plot based on weighted UniFrac distances between samples from ‘Cd-0_post’, ‘Cd-10_post’ and ‘Cd-50_post’ (rarefied feature-table; 30,000 sequences per sample). For the network analysis, we employed the default dissimilarity index (Jaccard, co-occurrence), with a maximum distance of 0.4 required to create an edge. To test whether the earthworm gut microbiota connectedness was higher within than between treatment groups, we extracted edges and nodes information from the network graph object using the R package ‘igraph’ (Csardi and Nepusz, 2006). We then produced a new network object using the ‘asNetwork’ function from the R package ‘intergraph’ (88) to apply an approximate maximum likelihood estimate based on a Monte Carlo scheme using the R package ‘ergm’ (89). We tested whether a model including nodes (factor ‘treatment group’) in addition to ‘edges’ explained the variation better than a model with only ‘edges’ using a chi-squared test.

To investigate those bacterial SVs that differed significantly between earthworms from the control and Cd-polluted groups, we applied ‘DESeq2’ (90) within ‘phyloseq’ and we further tested whether the SV read abundances differed between treatment groups at the genus and phylum level by using ANOVA and the Post-Hoc Tukey HSD test.

Finally, we tested those bacterial taxa from earthworm microbiotas that were significantly attributable to ‘Cd-0_post’, ‘Cd-10_post’ or ‘Cd-50_post’ by applying an ‘indicator species analysis’ with 999 permutations (R package ‘indicspecies’ (54)). This method enabled us to analyse the relative abundance and occurrence of SVs in samples of the various treatment groups in order to identify the SVs that significantly characterized the respective groups. The maximal index of 1 indicated that a certain SV was present in all individuals of the group of interest. An indicator value less than 1 meant that the SV was either present in more than one group or that it was not present in all members of this group. This method has been previously suggested for the definition of biomarkers of exposure to chemicals (35).

## Acknowledgements

We are grateful to Kerstin Wilhelm for laboratory guidance of M.S. during her visit at the Institute for Evolutionary Ecology and Conservation Genomics at the University of Ulm and Victoria Drechsel from the University of Innsbruck, for help with earthworm maintenance and feeding. S.S thanks the Ministry of Science, Research and Art, Baden-Württemberg for start-up funding of the EcoHealth initiative ‘Microbial Drivers in Plant and Animal Health’.

## Author contribution

M.S., M.H., S.M., and S.S. conceived the study. M.S. developed the theory and performed the laboratory work. S.M. performed bioinformatic analyses. M.S. and S.M. carried out the statistical analyses. M.S. and S.M. wrote the manuscript. M.H. and S.S. critically reviewed the manuscript. The authors declare that there is no conflict of interest.

## References

1. Järup L, Akesson A. 2009. Current status of cadmium as an environmental health problem. Toxicol Appl Pharmacol 238:201–8.

2. Jin Y, Wu S, Zeng Z, Fu Z. 2017. Effects of environmental pollutants on gut microbiota. Environ Pollut 222:1–9.

3. Gremion F, Chatzinotas A, Harms H. 2003. Comparative 16S rDNA and 16S rRNA sequence analysis indicates that Actinobacteria might be a dominant part of the metabolically active bacteria in heavy metal-contaminated bulk and rhizosphere soil. Environ Microbiol 5:896–907.

4. Bamborough L, Cummings SP. 2009. The impact of zinc and lead concentrations and seasonal variation on bacterial and actinobacterial community structure in a metallophytic grassland soil. Folia Microbiol (Praha) 54:327–34.

5. Singh BK, Quince C, Macdonald CA, Khachane A, Thomas N, Al-Soud WA, Sørensen SJ, He Z, White D, Sinclair A, Crooks B, Zhou J, Campbell CD. 2014. Loss of microbial diversity in soils is coincident with reductions in some specialized functions. Environ Microbiol 16:2408–20.

6. Chen YP, Liu Q, Liu YJ, Jia FA, He XH. 2014. Responses of soil microbial activity to cadmium pollution and elevated CO2. Sci Rep 4:4287.

7. Chodak M, Gołębiewski M, Morawska-Płoskonka J, Kuduk K, Niklińska M. 2013. Diversity of microorganisms from forest soils differently polluted with heavy metals. Appl Soil Ecol 64:7–14.

8. Frostegård A, Tunlid A, Bååth E. 1993. Phospholipid Fatty Acid Composition, Biomass, and Activity of Microbial Communities from Two Soil Types Experimentally Exposed to Different Heavy Metals. Appl Environ Microbiol 59:3605–17.

9. Gołębiewski M, Deja-Sikora E, Cichosz M, Tretyn A, Wróbel B. 2014. 16S rDNA pyrosequencing analysis of bacterial community in heavy metals polluted soils. Microb Ecol 67:635–47.

10. Sheik CS, Mitchell TW, Rizvi FZ, Rehman Y, Faisal M, Hasnain S, McInerney MJ, Krumholz LR. 2012. Exposure of Soil Microbial Communities to Chromium and Arsenic Alters Their Diversity and Structure. PLoS One 7:e40059.

11. Gans J, Wolinsky M, Dunbar J. 2005. Computational improvements reveal great bacterial diversity and high metal toxicity in soil. Science 309:1387–90.

12. Liu D, Lian B, Wang B, Jiang G. 2011. Degradation of potassium rock by earthworms and responses of bacterial communities in its gut and surrounding substrates after being fed with mineral. PLoS One 6:e28803.

13. Brito-Vega H, Espinosa-Victoria D. 2009. Bacterial diversity in the digestive tract of earthworms (Oligochaeta). J Biol Sci 9:192–199.

14. Pass DA, Morgan AJ, Read DS, Field D, Weightman AJ, Kille P. 2015. The effect of anthropogenic arsenic contamination on the earthworm microbiome. Environ Microbiol 17:1884–96.

15. Nechitaylo TY, Yakimov MM, Godinho M, Timmis KN, Belogolova E, Byzov BA, Kurakov A V, Jones DL, Golyshin PN, Belogolova E, Golyshin PN, Yakimov MM, Byzov BA, Kurakov A V, Timmis KN, Jones DL. 2010. Effect of the Earthworms Lumbricus terrestris and Aporrectodea caliginosa on Bacterial Diversity in Soil. Microb Ecol 59:574–587.

16. Aira M, Bybee S, Pérez-Losada M, Domínguez J. 2015. Feeding on microbiomes: effects of detritivory on the taxonomic and phylogenetic bacterial composition of animal manures. FEMS Microbiol Ecol 91:1–10.

17. Drake HL, Horn MA. 2007. As the Worm Turns: The Earthworm Gut as a Transient Habitat for Soil Microbial Biomes. Annu Rev Microbiol 61:169–189.

18. Knapp B a., Podmirseg SM, Seeber J, Meyer E, Insam H. 2009. Diet-related composition of the gut microbiota of Lumbricus rubellus as revealed by a molecular fingerprinting technique and cloning. Soil Biol Biochem 41:2299–2307.

19. Khomyakov N V., Kharin SA, Nechitailo TY, Golyshin PN, Kurakov A V., Byzov BA, Zvyagintsev DG. 2007. Reaction of microorganisms to the digestive fluid of earthworms. Microbiology 76:45–54.

20. Dishaw LJ, Cannon JP, Litman GW, Parker W. 2014. Immune-directed support of rich microbial communities in the gut has ancient roots. Dev Comp Immunol 47:36–51.

21. Parthasarathi K, Ranganathan LS, Anandi V, Zeyer J. 2007. Diversity of microflora in the gut and casts of tropical composting earthworms reared on different substrates. J Environ Biol 28:87–97.

22. Idowu AB, Edema MO, Adeyi AO. 2008. Gut Microflora and Microfauna of Earthworm Species in the Soils of the Research Farms of the University of Agriculture, Abeokuta, Nigeria. Biol Agric Hortic 25:185–200.

23. Singh A, Singh DP, Tiwari R, Kumar K, Singh RV, Singh S, Prasanna R, Saxena AK, Nain L. 2015. Taxonomic and functional annotation of gut bacterial communities of Eisenia foetida and Perionyx excavatus. Microbiol Res 175:48–56.

24. Zhang W, Guo R, Yang Y, Ding J, Zhang Y. 2016. Long-term effect of heavy-metal pollution on diversity of gastrointestinal microbial community of Bufo raddei. Toxicol Lett 258:192–197.

25. Breton J, Daniel C, Dewulf J, Pothion S, Froux N, Sauty M, Thomas P, Pot B, Foligné B. 2013. Gut microbiota limits heavy metals burden caused by chronic oral exposure. Toxicol Lett 222:132–138.

26. Breton J, Massart S, Vandamme P, De Brandt E, Pot B, Foligné B. 2013. Ecotoxicology inside the gut: impact of heavy metals on the mouse microbiome. BMC Pharmacol Toxicol 14:1–11.

27. Liu Y, Li Y, Liu K, Shen J. 2014. Exposing to cadmium stress cause profound toxic effect on microbiota of the mice intestinal tract. PLoS One 9:e85323.

28. Zhang S, Jin Y, Zeng Z, Liu Z, Fu Z. 2015. Subchronic Exposure of Mice to Cadmium Perturbs Their Hepatic Energy Metabolism and Gut Microbiome. Chem Res Toxicol 28:2000–9.

29. Zhai Q, Yu L, Li T, Zhu J, Zhang C, Zhao J, Zhang H, Chen W. 2017. Effect of dietary probiotic supplementation on intestinal microbiota and physiological conditions of Nile tilapia (Oreochromis niloticus) under waterborne cadmium exposure. Antonie Van Leeuwenhoek 110:501–513.

30. Wu B, Cui H, Peng X, Pan K, Fang J, Zuo Z, Deng J, Wang X, Huang J. 2014. Toxicological effects of dietary nickel chloride on intestinal microbiota. Ecotoxicol Environ Saf 109:70–6.

31. Guo X, Liu S, Wang Z, Zhang X, Li M, Wu B. 2014. Metagenomic profiles and antibiotic resistance genes in gut microbiota of mice exposed to arsenic and iron. Chemosphere 112:1–8.

32. McPhee JB, Schertzer JD. 2015. Immunometabolism of obesity and diabetes: microbiota link compartmentalized immunity in the gut to metabolic tissue inflammation. Clin Sci (Lond) 129:1083–1096.

33. Round JL, Mazmanian SK. 2009. The gut microbiota shapes intestinal immune responses during health and disease. Nat Rev Immunol 9:313–23.

34. Palm NW, de Zoete MR, Flavell R a. 2015. Immune-microbiota interactions in health and disease. Clin Immunol 159:122–127.

35. Gaulke CA, Barton CL, Proffitt S, Tanguay RL, Sharpton TJ. 2016. Triclosan exposure is associated with rapid restructuring of the microbiome in adult zebrafish. PLoS One 11:1–20.

36. Gómez-Brandón M, Aira M, Lores M, Domínguez J. 2011. Epigeic Earthworms Exert a Bottleneck Effect on Microbial Communities through Gut Associated Processes. PLoS One 6:e24786.

37. Aira M, Domínguez J. 2014. Changes in nutrient pools, microbial biomass and microbial activity in soils after transit through the gut of three endogeic earthworm species of the genus Postandrilus Qui and Bouché, 1998. J Soils Sediments 14:1335–1340.

38. Furlong MA, Singleton DR, Coleman DC, Whitman WB. 2002. Molecular and culture-based analyses of prokaryotic communities from an agricultural soil and the burrows and casts of the earthworm Lumbricus rubellus. Appl Environ Microbiol 68:1265–79.

39. Schönholzer F, Hahn D, Zarda B, Zeyer J. 2002. Automated image analysis and in situ hybridization as tools to study bacterial populations in food resources, gut and cast of Lumbricus terrestris L. J Microbiol Methods 48:53–68.

40. Nechitaylo TY, Timmis KN, Golyshin PN. 2009. “Candidatus Lumbricincola”, a novel lineage of uncultured Mollicutes from earthworms of family Lumbricidae. Environ Microbiol 11:1016–26.

41. Ba Q, Li M, Chen P, Huang C, Duan X, Lu L, Li J, Chu R, Xie D, Song H, Wu Y, Ying H, Jia X, Wang H. 2016. Gender-Dependent Effects of Cadmium Exposure in Early Life on Gut Microbiota and Fat Accumulation in Mice. Environ Health Perspect 125:1–5.

42. Ellis RJ, Morgan P, Weightman AJ, Fry JC. 2003. Cultivation-dependent and -independent approaches for determining bacterial diversity in heavy-metal-contaminated soil. Appl Environ Microbiol 69:3223–3230.

43. Sułowicz S, Płociniczak T, Piotrowska-Seget Z, Kozdrój J. 2011. Significance of silver birch and bushgrass for establishment of microbial heterotrophic community in a metal-mine spoil heap. Water Air Soil Pollut 214:205–218.

44. Evtushenko LI, Ariskina E V. 2012. Bergey’s Manual of Systematic Bacteriology, Volume 5: The Actinobacteria, Family II: Nocardioidaceae. Springer-Verlag New York.

45. Sani R, Peyton B, Smith W, Apel W, Petersen J. 2002. Dissimilatory reduction of Cr(VI), Fe(III), and U(VI) by Cellulomonas isolates. Appl Microbiol Biotechnol 60:192–199.

46. Hegazi RM, El-Gendy NS, El-Feky AA, Moustafa YM, El-Ezbewy S, El-Gemaee GH. 2007. Impact of heavy metals on biodegradation of phenanthrene by Cellulomonas hominis strain N2. J Pure Appl Microbiol 1:165–175.

47. Govarthanan M, Mythili R, Selvankumar T, Kamala-Kannan S, Rajasekar A, Chang Y-C. 2016. Bioremediation of heavy metals using an endophytic bacterium Paenibacillus sp. RM isolated from the roots of Tridax procumbens. 3 Biotech 6:242.

48. Rawat M, Rai JPN. 2012. Adsorption of Heavy Metals by *Paenibacillus validus* Strain MP5 Isolated from Industrial Effluent–Polluted Soil. Bioremediat J 16:66–73.

49. Rajbanshi A. 2009. Study on Heavy Metal Resistant Bacteria in Guheswori Sewage Treatment Plant. Our Nat 6:52–57.

50. Benmalek Y, Halouane A, Hacene H, Fardeau ML. 2014. Resistance to heavy metals and bioaccumulation of lead and zinc by Chryseobacterium solincola strain 1YB-R12T isolated from soil. Int J Environ Eng 6:68.

51. Mosa KA, Saadoun I, Kumar K, Helmy M, Dhankher OP. 2016. Potential Biotechnological Strategies for the Cleanup of Heavy Metals and Metalloids. Front Plant Sci 7:303.

52. Xie P, Hao X, Herzberg M, Luo Y, Nies DH, Wei G. 2015. Genomic analyses of metal resistance genes in three plant growth promoting bacteria of legume plants in Northwest mine tailings, China. J Environ Sci 27:179–187.

53. Zhang M, Huang F, Wang G, Liu X, Wen J, Zhang X, Huang Y, Xia Y. 2017. Geographic distribution of cadmium and its interaction with the microbial community in the Longjiang River: risk evaluation after a shocking pollution accident. Sci Rep 7:227.

54. De Cáceres M, Legendre P. 2009. Associations between species and groups of sites: indices and statistical inference. Ecology 90:3566–74.

55. Li F, Hullar MAJ, Schwarz Y, Lampe JW. 2009. Human Gut Bacterial Communities Are Altered by Addition of Cruciferous Vegetables to a Controlled Fruit- and Vegetable-Free Diet 1–3. J Nutr 139:1685–1691.

56. Galimanas V, Hall M, Singh N, Lynch MD, Goldberg M, Tenenbaum H, Cvitkovitch D, Neufeld J, Senadheera D. 2014. Bacterial community composition of chronic periodontitis and novel oral sampling sites for detecting disease indicators. Microbiome 2:32.

57. Kembel SW, Jones E, Kline J, Northcutt D, Stenson J, Womack AM, Bohannan BJ, Brown GZ, Green JL. 2012. Architectural design influences the diversity and structure of the built environment microbiome. ISME J 6:1469–1479.

58. Horn MA, Ihssen J, Matthies C, Schramm A, Acker G, Drake HL. 2005. Dechloromonas denitrificans sp. nov., Flavobacterium denitrificans sp. nov., Paenibacillus anaericanus sp. nov. and Paenibacillus terrae strain MH72, N2O-producing bacteria isolated from the gut of the earthworm Aporrectodea caliginosa. Int J Syst Evol Microbiol 55:1255–1265.

59. Kumar V, Bharti A, Negi YK, Gusain O, Pandey P, Bisht GS. 2012. Screening of actinomycetes from earthworm castings for their antimicrobial activity and industrial enzymes. Braz J Microbiol 43:205–14.

60. Polyanskaya LM, Babkina NI, Zenova GM, Zvyagintsev DG. 1996. Fate of actinomycetes in the intestinal tract of soil invertebrates fed on streptomycete spores. Microbiol New York 65:493–498.

61. Park J, Lee PA, Lee H-H, Choi K, Lee S-W, Seo Y-S. 2017. Comparative Genome Analysis of Rathayibacter tritici NCPPB 1953 with Rathayibacter toxicus Strains Can Facilitate Studies on Mechanisms of Nematode Association and Host Infection. plant Pathol J 33:370–381.

62. Aira M, Olcina J, Pérez-Losada M, Domínguez J. 2016. Characterization of the bacterial communities of casts from Eisenia andrei fed with different substrates. Appl Soil Ecol 98:103–111.

63. Kim H-J, Shin K-H, Cha C-J, Hur H-G. 2004. Analysis of Aerobic and Culturable Bacterial Community Structures in Earthworm (Eisenia fetida) Intestine. Agric Chem Biotechnol 47:137–142.

64. Verma K, Agrawal N, Farooq M, Misra RB, Hans RK. 2006. Endosulfan degradation by a Rhodococcus strain isolated from earthworm gut. Ecotoxicol Environ Saf 64:377–381.

65. Joshi A, Jaiswal P. 2013. Micro organisms living in zinc contaminated soil -a review. IOSR J Pharm Biol Sci 6:67–72.

66. Corretto E, Antonielli L, Sessitsch A, Compant S, Höfer C, Puschenreiter M, Brader G. 2017. Complete genome sequence of the heavy metal resistant bacterium Agromyces aureus AR33T and comparison with related Actinobacteria. Stand Genomic Sci 12:2.

67. Vela-Cano M, Castellano-Hinojosa A, Vivas AF, Victoria M, Toledo M. 2014. Effect of Heavy Metals on the Growth of Bacteria Isolated from Sewage Sludge Compost Tea. Adv Microbiol 4:644–655.

68. Goswami L, Arul Manikandan N, Pakshirajan K, Pugazhenthi G. 2017. Simultaneous heavy metal removal and anthracene biodegradation by the oleaginous bacteria Rhodococcus opacus. 3 Biotech 7:37.

69. Anwar M, Ali S, Asaad AT. 2017. Evaluation of Heavy Metals Resistant Micrococcus sp. Isolated from Rivers in Basra, Iraq. J Bioremediation Biodegrad 8:1–4.

70. Montero-Calasanz M del C, Hofner B, Göker M, Rohde M, Spröer C, Hezbri K, Gtari M, Schumann P, Klenk H-P. 2014. Geodermatophilus poikilotrophi sp. nov.: a multitolerant actinomycete isolated from dolomitic marble. Biomed Res Int 2014:914767.

71. Kant R, van Passel MWJ, Palva A, Lucas S, Lapidus A, Glavina del Rio T, Dalin E, Tice H, Bruce D, Goodwin L, Pitluck S, Larimer FW, Land ML, Hauser L, Sangwan P, de Vos WM, Janssen PH, Smidt H. 2011. Genome sequence of Chthoniobacter flavus Ellin428, an aerobic heterotrophic soil bacterium. J Bacteriol 193:2902–3.

72. Panke-Buisse K, Poole AC, Goodrich JK, Ley RE, Kao-Kniffin J. 2014. Selection on soil microbiomes reveals reproducible impacts on plant function. ISME J 9:980–989.

73. Vishnivetskaya TA, Mosher JJ, Palumbo A V, Yang ZK, Podar M, Brown SD, Brooks SC, Gu B, Southworth GR, Drake MM, Brandt CC, Elias DA. 2011. Mercury and other heavy metals influence bacterial community structure in contaminated Tennessee streams. Appl Environ Microbiol 77:302–11.

74. Byzov BA, Nechitaylo TY, Bumazhkin BK, Kurakov A V., Golyshin PN, Zvyagintsev DG. 2009. Culturable microorganisms from the earthworm digestive tract. Microbiology 78:360–368.

75. Drewniak L, Krawczyk PS, Mielnicki S, Adamska D, Sobczak A, Lipinski L, Burec-Drewniak W, Sklodowska A. 2016. Physiological and Metagenomic Analyses of Microbial Mats Involved in Self-Purification of Mine Waters Contaminated with Heavy Metals. Front Microbiol 7:1252.

76. Caporaso JG, Lauber CL, Walters WA, Berg-Lyons D, Huntley J, Fierer N, Owens SM, Betley J, Fraser L, Bauer M, Gormley N, Gilbert JA, Smith G, Knight R. 2012. Ultra-high-throughput microbial community analysis on the Illumina HiSeq and MiSeq platforms. ISME J 6:1621–1624.

77. Kuczynski J, Lauber CL, Walters WA, Parfrey LW, Clemente JC, Gevers D, Knight R. 2011. Experimental and analytical tools for studying the human microbiome. Nat Rev Genet 13:47–58.

78. Caporaso JG, Kuczynski J, Stombaugh J, Bittinger K, Bushman FD, Costello EK, Fierer N, Peña AG, Goodrich JK, Gordon JI, Huttley GA, Kelley ST, Knights D, Koenig JE, Ley RE, Lozupone CA, McDonald D, Muegge BD, Pirrung M, Reeder J, Sevinsky JR, Turnbaugh PJ, Walters WA, Widmann J, Yatsunenko T, Zaneveld J, Knight R. 2010. QIIME allows analysis of high-throughput community sequencing data. Nat Methods 7:335–336.

79. Callahan BJ, McMurdie PJ, Rosen MJ, Han AW, Johnson AJA, Holmes SP. 2016. DADA2: High-resolution sample inference from Illumina amplicon data. Nat Methods 13:581–583.

80. Katoh K, Standley DM. 2013. MAFFT Multiple Sequence Alignment Software Version 7: Improvements in Performance and Usability. Mol Biol Evol 30:772–780.

81. Price MN, Dehal PS, Arkin AP. 2010. FastTree 2 –Approximately Maximum-Likelihood Trees for Large Alignments. PLoS One 5:e9490.

82. Faith DP. 1992. Conservation evaluation and phylogenetic diversity. Biol Conserv 61:1–10.

83. Lozupone CA, Hamady M, Kelley ST, Knight R. 2007. Quantitative and qualitative beta diversity measures lead to different insights into factors that structure microbial communities. Appl Environ Microbiol 73:1576–85.

84. R Development Core Team. 2011. R: A Language and Environment for Statistical Computing. Vienna, Austria: the R Foundation for Statistical Computing.

85. McDonald D, Clemente JC, Kuczynski J, Rideout JR, Stombaugh J, Wendel D, Wilke A, Huse S, Hufnagle J, Meyer F, Knight R, Caporaso JG. 2012. The Biological Observation Matrix (BIOM) format or: how I learned to stop worrying and love the ome-ome. Gigascience 1:7.

86. McMurdie PJ, Holmes S. 2013. phyloseq: An R Package for Reproducible Interactive Analysis and Graphics of Microbiome Census Data. PLoS One 8:e61217.

87. Csardi G, Nepusz T. 2006. The Igraph Software Package for Complex Network Research. InterJournal Complex Sy:1695.

88. Maintainer MB, Bojanowski M. 2016. Title Coercion Routines for Network Data Objects.

89. Hunter DR, Handcock MS, Butts CT, Goodreau SM, Morris M. 2008. ergm: A Package to Fit, Simulate and Diagnose Exponential-Family Models for Networks. JSS J Stat Softw 24.

90. Love MI, Huber W, Anders S. 2014. Moderated estimation of fold change and dispersion for RNA-seq data with DESeq2. Genome Biol 15:550.

